# Small molecule v-ATPase inhibitor Etidronate lowers levels of ALS protein ataxin-2

**DOI:** 10.1101/2021.12.20.473567

**Authors:** Garam Kim, Lisa Nakayama, Jacob A. Blum, Tetsuya Akiyama, Steven Boeynaems, Meena Chakraborty, Julien Couthouis, Eduardo Tassoni-Tsuchida, Caitlin M. Rodriguez, Michael C. Bassik, Aaron D. Gitler

## Abstract

Antisense oligonucleotide therapy targeting *ATXN2*—a gene in which mutations cause neurodegenerative diseases spinocerebellar ataxia type 2 and amyotrophic lateral sclerosis—has entered clinical trials in humans. Additional methods to lower ataxin-2 levels would be beneficial not only in uncovering potentially cheaper or less invasive therapies, but also in gaining greater mechanistic insight into how ataxin-2 is normally regulated. We performed a genome-wide fluorescence activated cell sorting (FACS)-based CRISPR screen in human cells and identified multiple subunits of the lysosomal vacuolar ATPase (v-ATPase) as regulators of ataxin-2 levels. We demonstrate that Etidronate—a U.S. Food and Drug Administration (FDA)-approved drug that inhibits the v-ATPase—lowers ataxin-2 protein levels in mouse and human neurons. Moreover, oral administration of the drug to mice in their water supply and food is sufficient to lower ataxin-2 levels in the brain. Thus, we uncover Etidronate as a safe and inexpensive compound for lowering ataxin-2 levels and demonstrate the utility of FACS-based screens for identifying targets to modulate levels of human disease proteins.

## Introduction

Many neurodegenerative diseases are associated with accumulation of misfolded proteins (Forman et al., 2004), and strategies to rid these proteins are emerging as a promising therapeutic approach (Bennett et al., 2021). TDP-43 is the disease protein in nearly all cases of amyotrophic lateral sclerosis (ALS) (Neumann et al., 2006). Despite the presence of TDP-43 pathology in ALS and numerous other neurodegenerative disorders (de Boer et al., 2020), targeting TDP-43 presents several challenges. TDP-43 is not only tightly regulated and its function essential in most cell types (Kim et al., 2020), but also reducing its levels *in vivo* results in embryonic lethality during development and motor phenotypes in adult mice (Kim et al., 2020; Yang et al., 2014).

Another approach could be to target a modifier of TDP-43 aggregation and toxicity. Ataxin-2—a polyglutamine (polyQ) protein for which long (>34) polyQ expansions cause spinocerebellar ataxia 2 (SCA2) (Imbert et al., 1996; Pulst et al., 1996; Sanpei et al., 1996) and intermediate-length (27-33) repeats are a risk factor for ALS (Elden et al., 2010)—is a potent modifier of TDP-43 toxicity (Armakola et al., 2011; Becker et al., 2017; Elden et al., 2010; Kim et al., 2014). Antisense oligonucleotides (ASOs) targeting ataxin-2 *in vivo* show marked protection against motor deficits and extend lifespan in TDP-43 overexpressing mice (Becker et al., 2017) and in SCA2 mice (Scoles et al., 2017). These results have motivated recent administration of ataxin-2 targeting ASOs to human ALS patients with or without polyQ expansions in a phase 1 clinical trial (ClinicalTrials.gov identifier: NCT04494256). Gene-based therapies, like ASOs, hold great promise to provide disease-modifying treatments for these devastating neurodegenerative diseases but are yet unproven for ALS patients. Despite this enormous progress, questions remain regarding the potential safety or dosage limitations of using ASOs, and little is known about how ataxin-2 is normally regulated. Thus, we searched for other ways to modulate ataxin-2 protein levels using an efficient genome-wide screening paradigm.

We developed a fluorescence activated cell sorting (FACS)-based screening method using pooled CRISPR/Cas9-mediated genome-wide deletion libraries in conjunction with antibody staining to detect endogenous protein levels. The screen revealed numerous novel genetic modifiers of ATXN2 protein levels, including multiple subunits of the lysosomal vacuolar ATPase (v-ATPase). We show that inhibiting the lysosomal v-ATPase with small molecule drugs (FDA-approved) results in decreased ataxin-2 protein levels in mouse and human iPSC-derived neurons, as well as *in vivo* in the brains of mice upon oral administration of the drug in their water supply and diet. These results demonstrate the potential to re-purpose a cheap, readily available small molecule drug as a novel treatment for ALS and SCA2 patients.

## Results

### FACS-based genome-wide CRISPR-Cas9 KO screens in human cells reveal modifiers of ataxin-2 protein levels

To find additional ways (e.g., new targets or pathways) to lower ataxin-2 levels and expand potential therapeutic strategies, we developed a fluorescence-activated cell sorting (FACS)-based genome-wide CRISPR knockout (KO) screen for modifiers of endogenous ataxin-2 protein levels (i.e., without protein tags like Flag or GFP or overexpression) (**Figure 1A**). We optimized conditions to sensitively detect changes in ataxin-2 protein levels in human cells by antibody staining and FACS (**Figure S1**) and engineered HeLa cells to stably express Cas9 along with either a blasticidin-resistance cassette (HeLa-Cas9-Blast) or blue fluorescent protein (BFP) (HeLa-Cas9-BFP). To create genome-wide KO cell lines, we transduced HeLa-Cas9-Blast and HeLa-Cas9-BFP cells (as biological replicates) with a lentiviral sgRNA library comprising 10 sgRNAs per gene—targeting ∼21,000 human genes—and ∼10,000 safe-targeting sgRNAs (Morgens et al., 2017) (**Figure 1A**). After fixing the cells in methanol and immunostaining them with antibodies targeting endogenous ataxin-2 and a control protein (GAPDH or β-actin), we used FACS to sort the highest and lowest 20% levels of endogenous ataxin-2 expressors relative to levels of the control protein (**Figure 1A**). We performed the screen four times, twice each with β-actin and GAPDH as the control proteins. We used two different control proteins instead of one to minimize the possibility of selecting hits that were simply global regulators of transcription, translation, or cell size, or otherwise idiosyncratic to the biology of a given control protein.

**Figure 1:**
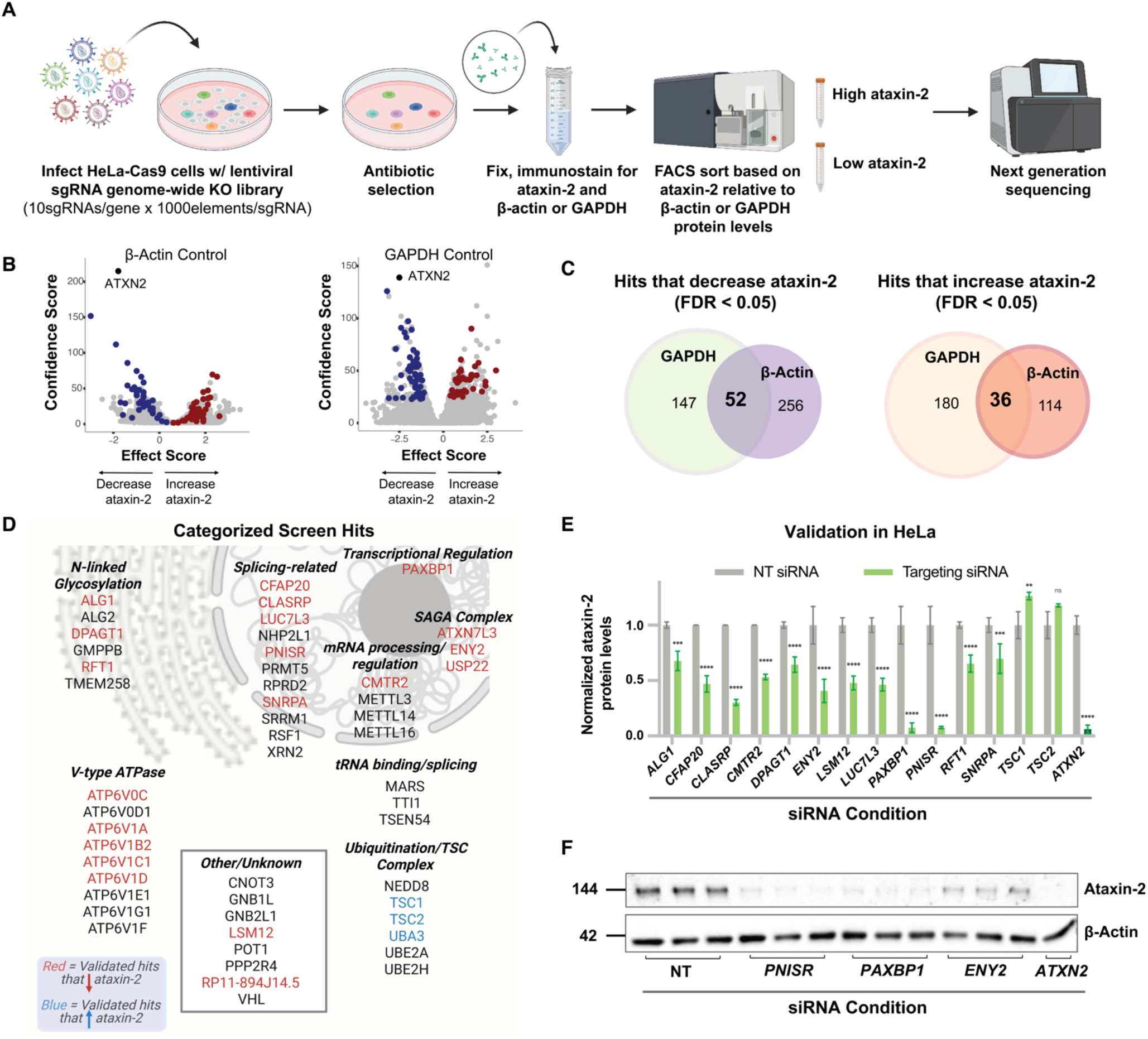
Genome-wide CRISPR–Cas9 KO screens in human cells identify regulators of ataxin-2 protein levels. **(A)** Pooled CRISPR–Cas9 screening paradigm. After transducing HeLa cells expressing Cas9 with a lentiviral sgRNA library, we fixed and co-immunostained the cells for ataxin-2 and a control protein (β-actin or GAPDH). We then used FACS to sort top and bottom 20% ataxin-2 expressors relative to control protein levels (duplicate sorts were performed using each control). After isolating genomic DNA from these populations, as well as the unsorted control population, we performed NGS to read sgRNA barcodes. **(B)** Volcano plots based on effect and confidence scores summarizing genes that modify ataxin-2 protein levels relative to β-actin (left) or GAPDH (right) levels when knocked out (FDR < 0.05). **(C)** Overlap between hits identified in β-actin-and GAPDH-controlled screens for hits that decrease (left) and increase (right) ataxin-2 protein levels when knocked out (FDR < 0.05). **(D)** Schematic of proteins encoded by selected hits (5% FDR), categorized by function and subcellular localization. **(E)** Validation of numerous top hit genes overlapping across all screens. We transfected HeLa-Cas9 cells with non-targeting (NT) siRNAs or with siRNAs targeting mRNA transcripts encoded by hit genes, then performed immunoblotting on lysates. Quantifications are normalized to the NT siRNA condition (mean±SD; analyzed using 2-way ANOVA; ****: p<0.0001, ***: p<0.001, **: p<0.01, *: p<0.05, ns: not significant). **(F)** Representative immunoblot of ataxin-2 and β-actin protein levels upon treatment with NT, *PNISR, PAXBP1*, and *ATXN2* siRNAs in HeLa cells.

After sorting cells based on ataxin-2 expression levels, we extracted genomic DNA and performed next generation sequencing (NGS) of the barcoded sgRNAs. Using the Cas9 high-throughput maximum-likelihood estimator (casTLE) algorithm (Morgens et al., 2016), we isolated genes that when knocked out increased or decreased ataxin-2 protein levels (**Figure 1B**). A false discovery rate (FDR) cutoff of 5% revealed an overlapping set of 52 gene knockouts that decreased—and 36 that increased—ataxin-2 levels across the four screens (**Figure 1C**). As expected, the strongest hit that lowered ataxin-2 levels was *ATXN2* itself, demonstrating the effectiveness of this screening approach (**Figure 1B**). The full list of hits from the screens is shown in Supplementary Data Table 1.

We individually validated screen hits by treating HeLa cells with siRNAs targeting those gene products followed by immunoblotting. We selected genes for follow-up based on a combination of significance (casTLE) score and effect size (Morgens et al., 2016) (**Supplementary Data Table 1**), as well as known and predicted direct (physical) and indirect (functional) associations with ataxin-2 or between hits. The list of hits included genes known to have a direct association with ataxin-2, such as *LSM12*, as well as many novel and potent genetic modifiers of ataxin-2 protein levels, such as *CFAP20, CLASRP, CMTR2, LUC7L3, PAXBP1, PNISR, ATP6V1A*, and *ATP6V1C1*, and others (**Figure 1D, 1E, and 2B**). Strikingly, some of the hits (e.g., *PNISR* and *PAXBP1*) lowered ataxin-2 levels as much as targeting ataxin-2 itself (**Figure 1F**). We further validated hits in the human neuroblastoma cell line (SH-SY5Y), where many— but not all—had a significant effect on ataxin-2 protein levels (**Figure S2A and S2B**). In addition to identifying numerous previously unknown regulators of ataxin-2, the high rate of validation of hits from the initial screen and its relative ease (a genome-wide screen can be completed in < 1 week from fixation to NGS analysis) suggests these FACS/antibody-staining based screens—using fixed cells to uncover regulators of protein levels—may be useful in many other contexts.

### Inhibition of lysosomal v-ATPase subunits via siRNA or small molecule drug treatment lowers ataxin-2 protein levels in vitro

We observed a striking number of genes encoding subunits of lysosomal vacuolar ATPases (v-ATPases) that decreased ataxin-2 levels when knocked out (**Figure 2A**). Lysosomal v-ATPases help to maintain an acidic pH (∼4.5) in the lysosome, by pumping protons from the cytosol to the lumen via consumption of ATP (Maxson and Grinstein, 2014). We used siRNAs and immunoblotting to confirm that knocking down numerous v-ATPase subunits leads to decreased ataxin-2 protein levels (**Figure 2B and 2C**). We also generated a constitutive knockout of the v-ATPase subunit ATP6V1A—a central subunit in v-ATPase function (Maxson and Grinstein, 2014)—using CRISPR-Cas9 in HeLa cells, which resulted in a marked decrease in ataxin-2 protein levels (**Figure 2E and 2F**). Levels of other polyQ disease proteins huntingtin and ataxin-1 were unaltered in these cells (**Figure S3**). To determine whether the mechanism of regulation was at the protein or RNA level, we performed RT-qPCR after siRNA treatments and observed no change in steady state *ATXN2* mRNA levels (**Figure 2D**). RNA-sequencing confirmed that knocking down a v-ATPase subunit using siRNAs did not lead to changes in *ATXN2* mRNA levels or other noteworthy transcriptional changes (**Figure S4**). These results provide evidence that the mechanism of v-ATPase regulation of ataxin-2 is at the protein level.

**Figure 2:**
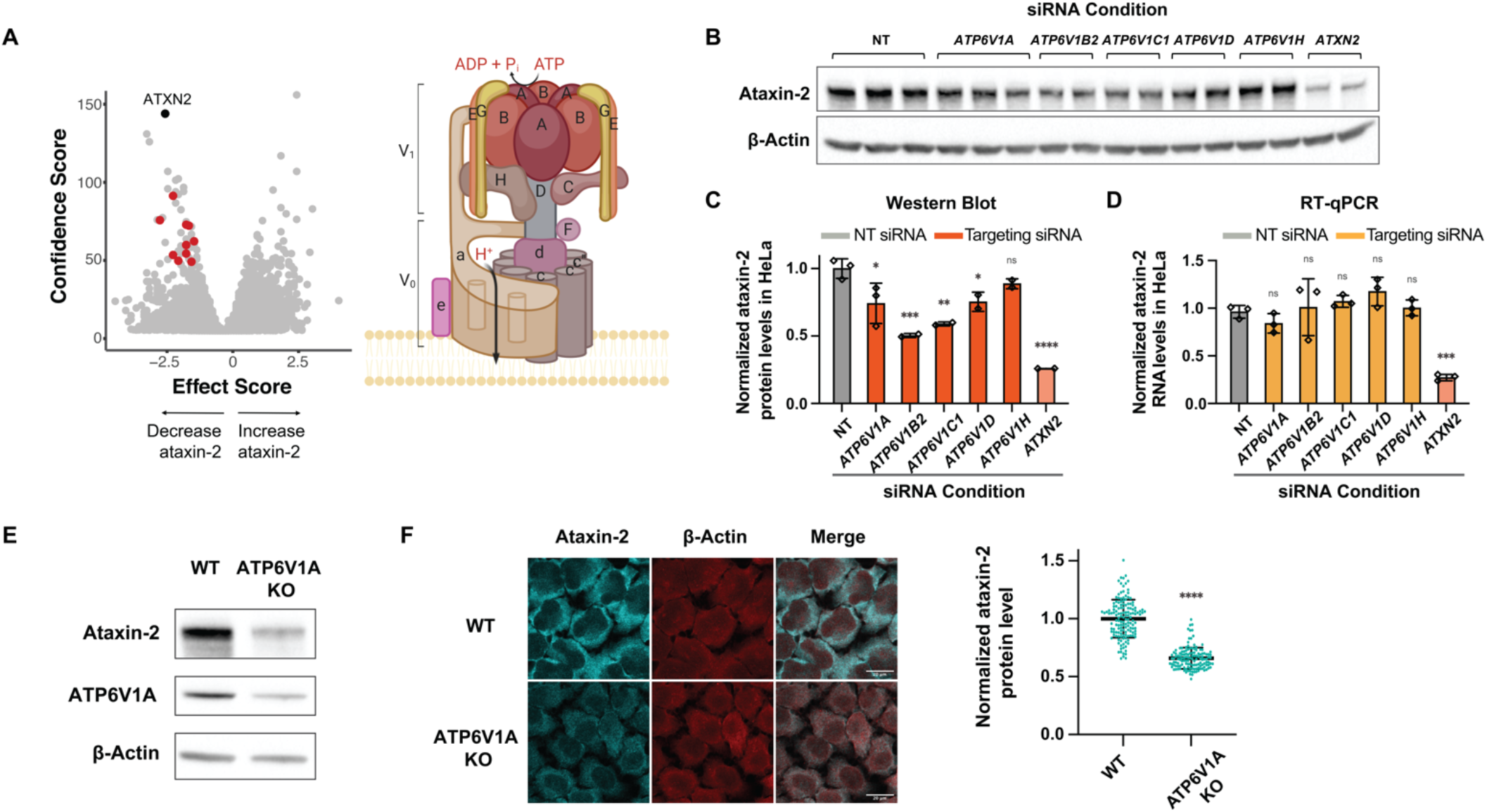
Inhibiting lysosomal v-ATPase leads to decreased ataxin-2 protein levels *in vitro*. **(A)** Left, volcano plot shows confidence score on y-axis and effect score on x-axis, with gene hits encoding subunits of lysosomal v-ATPases highlighted in red. Right, a representation of the lysosomal v-ATPase, with its V_0_ and V_1_ domains, as well as individual subunits. **(B)** Immunoblot and **(C)** Quantification of ataxin-2 protein levels after HeLa cells were transfected with siRNAs against various v-ATPase subunits. **(D)** RT-qPCR quantification of *ATXN2* RNA levels after siRNA knockdown of v-ATPase subunits in HeLa cells. Values normalized to β-actin RNA levels. Quantifications for c and d are normalized to the NT siRNA condition (mean ± SD; analyzed using one-way ANOVA with post-hoc Dunnett’s multiple comparisons tests; ****: p<0.0001, ***: p<0.001, **: p<0.01, *: p<0.05, ns: not significant). **(E)** Immunoblot on lysates from WT or *ATP6V1A* KO HeLa cell lines. **(F)** Representative microscopy images of WT or *ATP6V1A* KO cells, stained for ataxin-2 or β-actin (scale bar = 20 μm). Ataxin-2 fluorescence quantifications are shown on the right (lines denote mean ± SD; analyzed using unpaired t-test; ****: p<0.0001).

In addition to being enriched as screen hits, the v-ATPases stood out for another reason: the availability of small molecule inhibitors. Etidronate is a small molecule (∼206 Daltons) lysosomal v-ATPase inhibitor (**Figure 3A**) that was FDA-approved in 1977 as a drug to treat Paget disease of bone (Altman et al., 1973). Interestingly, Paget disease of bone has been connected to TDP-43 proteinopathy (Gitcho et al., 2009; Neumann et al., 2007; Watts et al., 2004). Etidronate is a bisphosphonate whose structural similarity to both ADP and phosphate is thought to be a mechanism by which it inhibits the v-ATPase (Altman et al., 1973). Etidronate has a 0.8901 predicted probability of crossing the blood brain barrier (BBB) according to ADMET *in silico* modeling (van de Waterbeemd and Gifford, 2003), and also a wide therapeutic index. Given these encouraging properties, especially its safety in humans, we tested whether Etidronate treatment could lower ataxin-2 levels in mouse primary neurons (**Figure 3B**). We also tested whether Thonzonium, another FDA-approved drug (∼511 Daltons) that inhibits the lysosomal v-ATPase through a different mechanism (uncoupling the proton transport and ATPase activity of the v-ATPase proton pumps), would decrease ataxin-2 protein levels.

**Figure 3:**
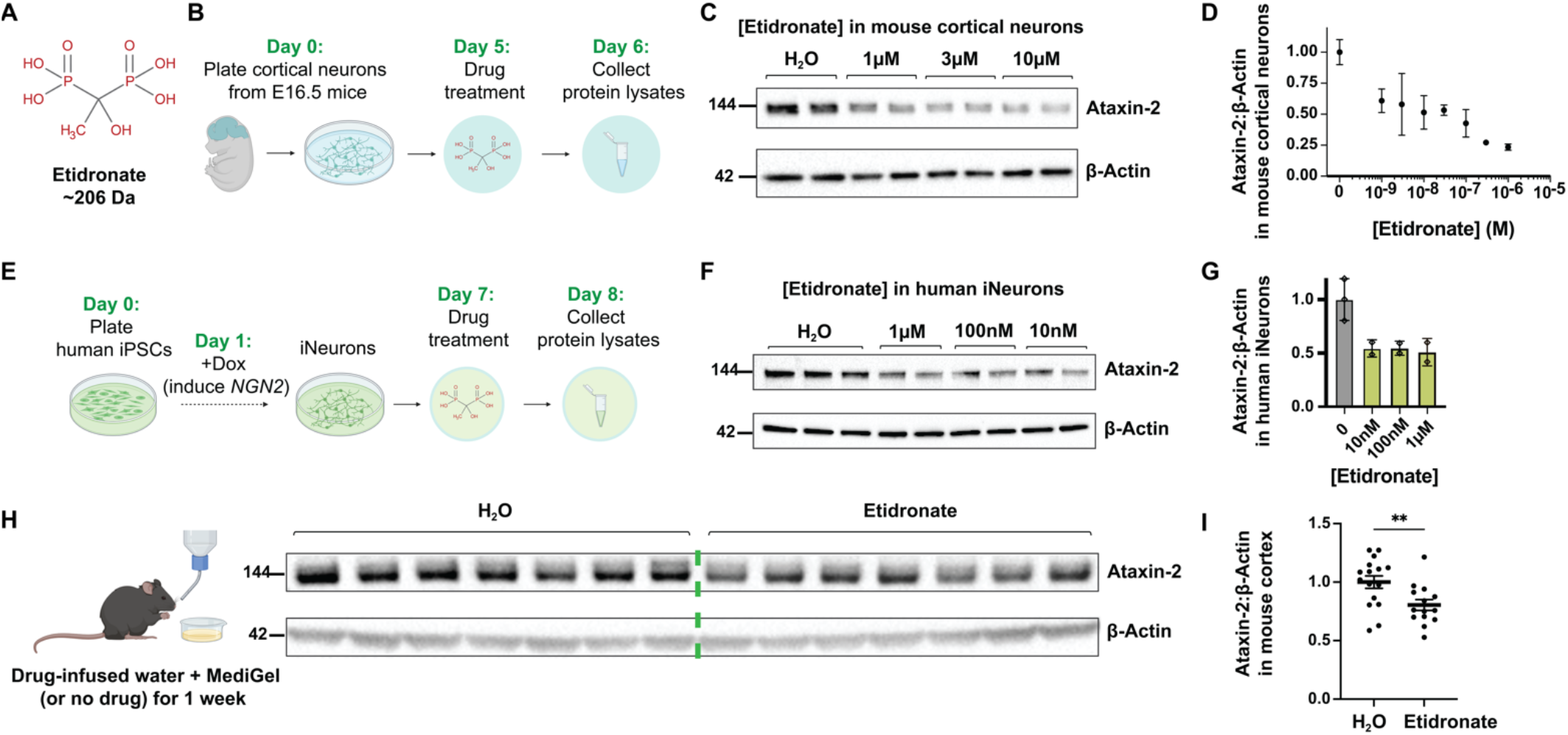
Small molecule drug Etidronate lowers ataxin-2 protein levels in mouse primary neurons, human iPSC-derived neurons, and *in vivo* in mice. **(A)** Structure of the small molecule drug Etidronate (C_2_H_8_O_7_P_2_). Etidronate is a bisphosphonate v-ATPase inhibitor. **(B)** Timeline of primary neuron plating from embryonic mouse cortex and drug treatment. **(C)** Immunoblot on lysates from mouse primary neurons treated with various doses of Etidronate. **(D)** Quantification of the dose-dependent effect of Etidronate on ataxin-2 (normalized to β-actin). **(E)** Timeline of induced neuron differentiation in a human iPSC line with *NGN2* stably integrated and drug treatment. **(F)** Immunoblot on lysates from human iPSC-derived neurons treated with 10 nM, 100 nM, or 1 μM of Etidronate. **(G)** Quantification of Figure 3F, with ataxin-2 protein levels normalized to β-actin (loading control) protein levels (mean ± SD). **(H)** Example immunoblot of cortex lysates from mice given normal or drug-infused water and MediGel®. **(I)** Quantification of immunoblots (e.g., Figure 3H) probing for ataxin-2 (normalized to β-actin protein levels) using lysates from cortices of mice that received water or drug treatment (mean ± SEM; analyzed using Welch’s t-test; **: p<0.01). We performed the experiment two independent times, for a total of n=15 in each treatment group.

Treatment of mouse cortical neurons with either drug for 24 hours resulted in a strong dose-dependent decrease in ataxin-2 protein levels (**Figure 3C, S5A, and S5B**). Although Thonzonium decreased ataxin-2 protein levels across a wide range of doses (**Figure S5A and S5B**), it became toxic to neurons at concentrations greater than 10 μM, as has been previously reported (Chan et al., 2012) (data not shown). However, Etidronate was not toxic across multiple orders of magnitude of concentrations (range of 10 nM to 100 μM tested) (**Figure 3D**). We also treated human iPSC-derived neurons with Etidronate (**Figure 3E**) and observed a similar safe and significant decrease in ataxin-2 levels (**Figure 3F and 3G**). Thus, two different v-ATPase inhibitors, working through distinct mechanisms, potently decrease ataxin-2 protein levels in neurons.

### Oral administration of Etidronate lowers ataxin-2 protein levels in vivo

We next tested if peripheral administration of Etidronate *in vivo* could decrease ataxin-2 levels in the brains of wild type (WT) mice. Given that oral administration of the drug mimics the most common mode of drug intake for humans and the wide range of doses that lowered ataxin-2 protein levels *in vitro* (**Figure 3C and 3D**), we dissolved Etidronate into the water supply (10 μM concentration) and MediGel® (ClearH_2_O) diet (30 μM concentration) of WT adult mice (∼3-4 months old) and allowed voluntary consumption over one week (**Figure 3H**). We performed immunoblotting on cortical extracts from mice that drank and ate either normal or drug-infused water and MediGel®, respectively. After a one-week treatment period, we observed a ∼20% decrease in ataxin-2 protein levels in the brains of mice that consumed Etidronate compared to the control group (n=15 in each group) (**Figure 3H and 3I**). These findings suggest that Etidronate administration in the water supply and MediGel® food is sufficient to lower levels of ataxin-2 in the brains of mice.

## Discussion

Here we used a highly efficient and robust FACS-based CRISPR screen to uncover numerous regulators of ataxin-2 levels—a validated target for ALS and SCA2 based on human genetics—including genes encoding several subunits of lysosomal v-ATPases for which small molecule drugs are available. One of these drugs, Etidronate, can safely and effectively decrease ataxin-2 levels in mouse and human neurons across a wide range of doses *in vitro*, as well as *in vivo* when orally administered to mice. Etidronate’s safety profile and molecular properties that indicate its ability to cross the BBB are highly favorable, providing a promising starting compound for future optimization to decrease ataxin-2 levels in the brain. Clinical trials in humans will be required to determine Etidronate’s safety and efficacy as a treatment for ALS or SCA2. ASO and gene therapy approaches show promise for neurodegenerative disease but are currently prohibitively expensive (hundreds of thousands of US dollars per year for ASOs or several million US dollars for a one-time gene therapy treatment). We purchased 1g of Etidronate for $50. The affordability of this compound could be especially useful in developing countries, like Cuba, where SCA2 is relatively common (Velázquez-Pérez et al., 2011). Etidronate has also been used to treat Paget disease of bone and osteoporosis for over forty years, providing the additional advantage of familiarity and experience with its application in the real world. Since many human diseases are caused by moderate increases or decreases in a gene product, we suspect that the protein levels-based screening approach shown here and in the accompanying manuscript by Rodriguez et al. (please see accompanying manuscript), as well as by others (Lu et al., 2013; Park et al., 2013; Rousseaux et al., 2018), could be broadly applicable to different human disease situations, including haploinsufficiency.

### Limitations of the study

Although we demonstrate that Etidronate can lower levels of ataxin-2 *in vitro* and *in vivo*, a limitation of this study is that we did not test whether the drug can rescue degeneration and motor phenotypes in a mouse model. Our previous study demonstrated that lowering levels of ataxin-2 prolonged survival and ameliorated motor impairments in a mouse model of TDP-43 proteinopathy (Becker et al., 2017). We faced difficulty in testing oral administration of Etidronate in this mouse model (Wils et al., 2010) because of the aggressive progression of disease (mice succumb prior to weaning age, at which point mice begin to eat/drink from their own food/water supplies). While future efforts will test the efficacy of this drug in ameliorating ALS phenotypes in other mouse models of ALS with later disease onset (Mitchell et al., 2015), we remain encouraged by the ability of Etidronate to lower ataxin-2 levels *in vivo*, as *ATXN2* is a previously validated therapeutic target in mouse models of ALS (Becker et al., 2017) and SCA2 (Scoles et al., 2017) as well as by human genetics (Elden et al., 2010; Scoles and Pulst, 2018).

## Acknowledgments

This work was supported by NIH grant R35NS097263 (A.D.G.), the Robert Packard Center for ALS Research at Johns Hopkins (A.D.G.), and the Brain Rejuvenation Project of the Stanford Neurosciences Institute (A.D.G.). G.K. is supported by a fellowship from the Stanford Knight-Hennessy Scholars Program. C.M.R. is supported by NIH grant F32NS116208-02. T.A. is supported by NIH grant 2T32AG047126-06A1 and a fellowship from the Takeda Science Foundation. Cell sorting/flow cytometry analysis for this project was completed on instruments in the Stanford Shared FACS Facility (SSFF), with a special thanks to Bianca Gomez. This work used the Genome Sequencing Service Center by Stanford Center for Genomics and Personalized Medicine Sequencing Center, supported by the grant award NIH S10OD025212, and NIH/NIDDK P30DK116074. Some of the computing for this project was performed on the Sherlock cluster. We would like to thank Stanford University and the Stanford Research Computing Center for providing computational resources and support that contributed to these research results.

## Figures

**Figure S1:**
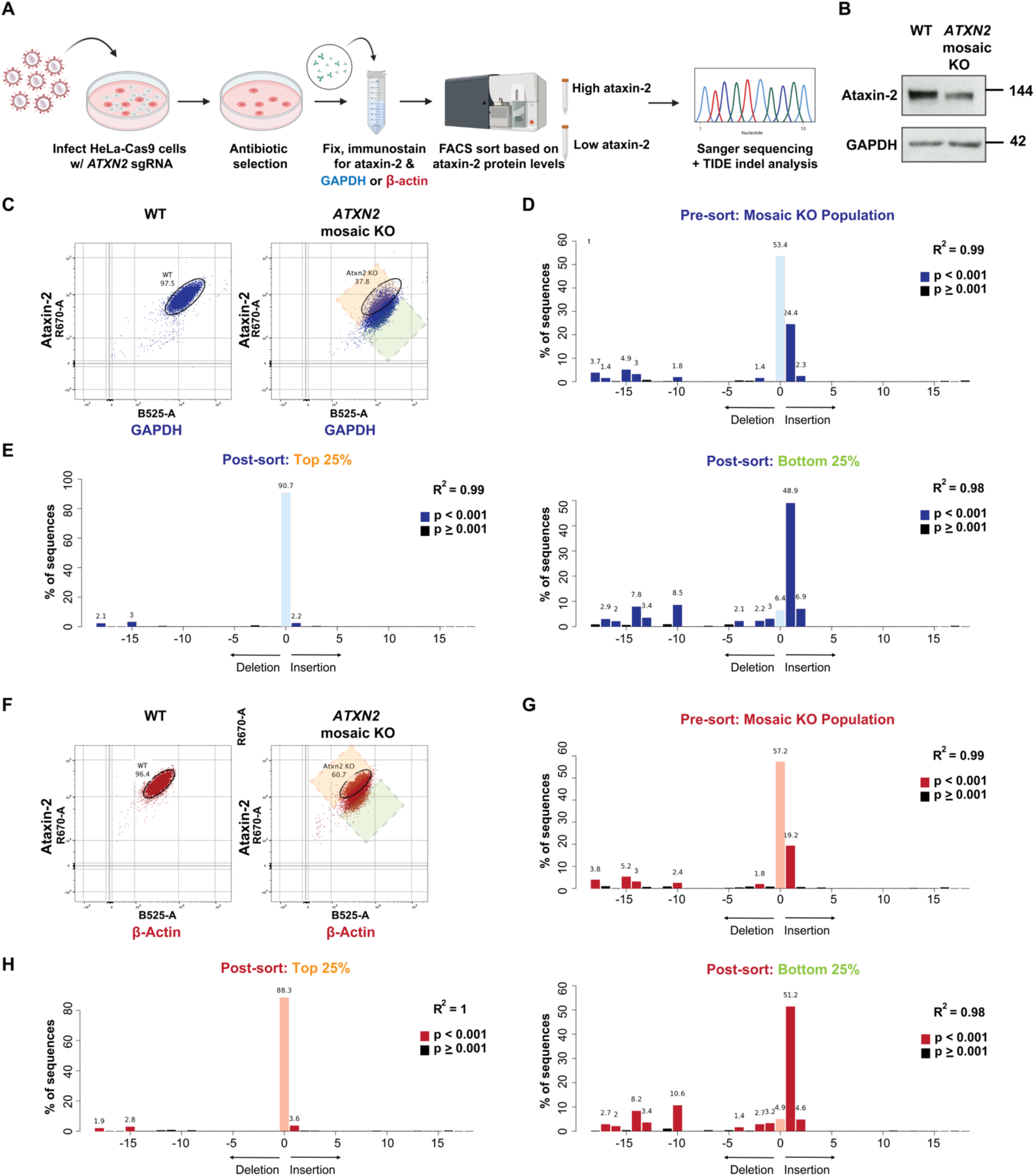
Calibration steps prior to conducting genome-wide screens. **(A)** Overview of screen optimization strategy. Briefly, HeLa cells expressing Cas9 were infected with a lentiviral sgRNA targeting ataxin-2, and puromycin was used to select for cells that received a guide. Cells were kept pooled to retain a mosaic population. These cells were then fixed in methanol, immunostained, and sorted using FACS for the top and bottom 25% of ataxin-2 expressors relative to a control protein (GAPDH or β-actin). The sorted and unsorted populations were Sanger sequenced and analyzed for insertions or deletions (indels) at the *ATXN2* locus. **(B)** Immunoblot of the WT and *ATXN2* mosaic KO populations, as generated in panel a. **(C)** Gating strategy for FACS. Left, FACS plot for WT population around which a gate is drawn; Right, mosaic population relative to the WT population. The green and orange gates represent the bottom and top 25% of *ATXN2* expressors relative to GAPDH, respectively. **(D)** Indel (insertion and deletion) analysis of the unsorted mosaic population (i.e. FACS plot in panel c, Right) when Sanger sequenced at the *ATXN2* locus, showing a mixture of cells containing various indels. **(E)** Indel analysis at *ATXN2* locus for the sorted populations. Left, overrepresentation of ATXN2 WT cells in the top 25% population (orange gate in panel c); Right, underrepresentation of WT cells / mostly various indels in the bottom 25% sorted population (green gate in panel c). **(F)** Same as in panel C, except using β-actin as control. **(G)** Same as in panel D, except using β-actin as control. **(H)** Same as in panel E, except using β-actin as control.

**Figure S2:**
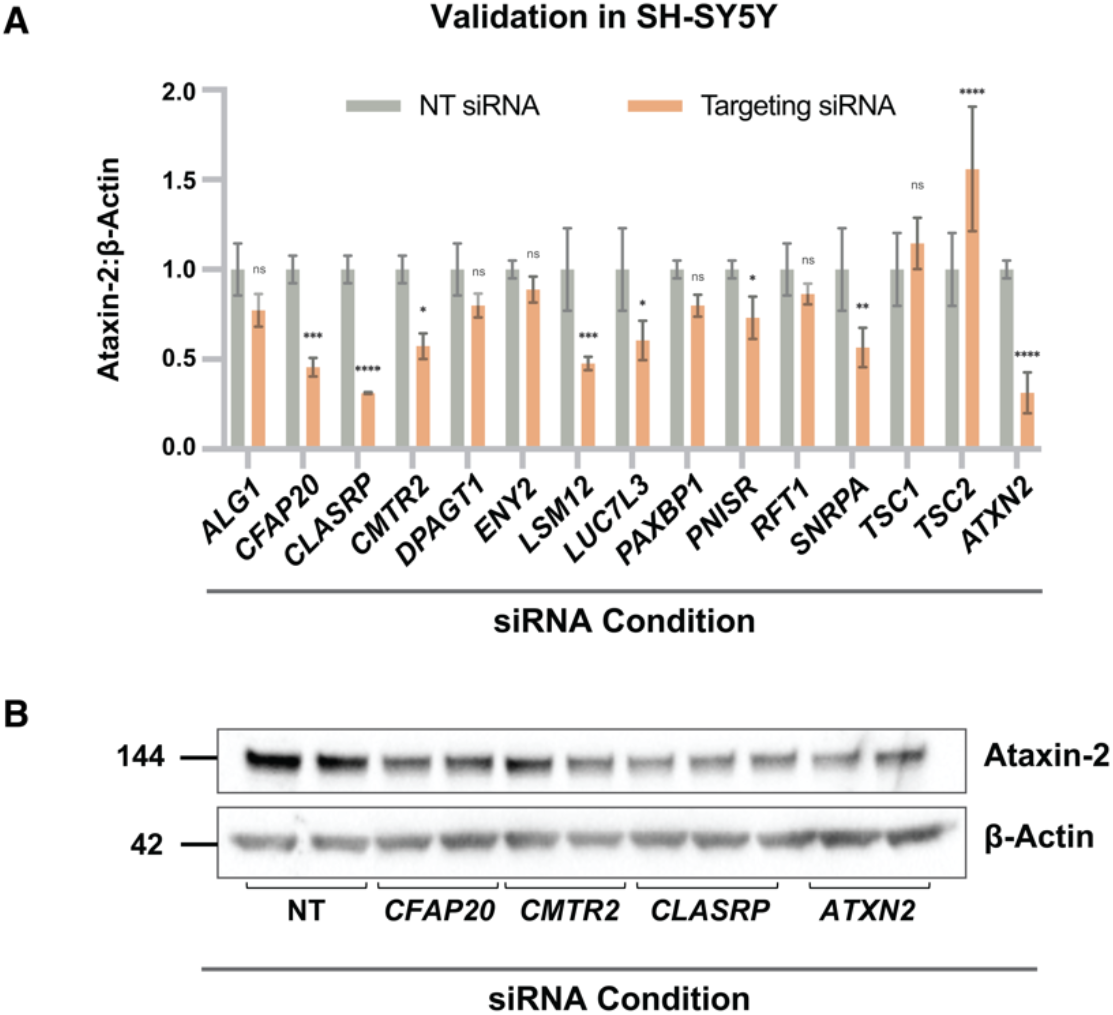
Validation of screen results in neuroblastoma cell line SH-SY5Y. **(A)** Validation of numerous top hit genes in SH-SY5Y cells using siRNA transfections and immunoblot analyses as in Fig. 1 e and f. Quantifications are normalized to the NT siRNA condition (mean ± SD; analyzed using 2-way ANOVA; ****: p<0.0001, ***: p<0.001, **: p<0.01, *: p<0.05, ns: not significant). **(B)** Representative immunoblot of ataxin-2 and β-actin protein levels upon application of NT, *CFAP20, CMTR2, CLASRP*, and *ATXN2* siRNAs to SH-SY5Y cells.

**Figure S3:**
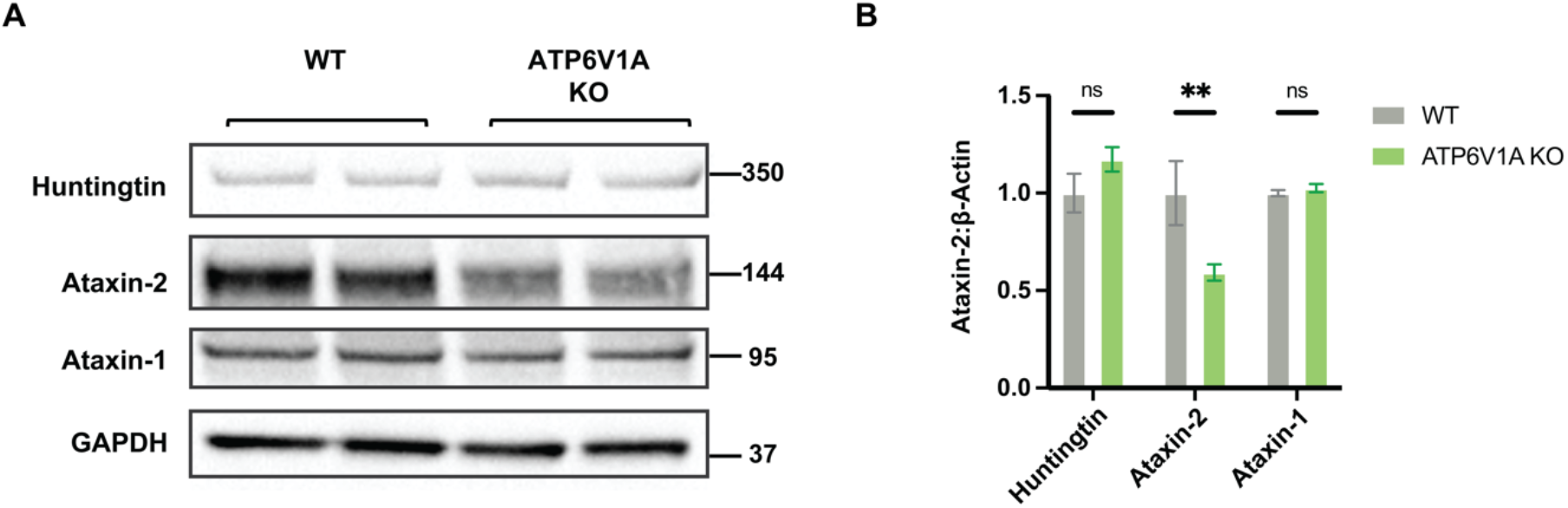
Other polyQ protein levels are unaltered in ATP6V1A KO cells. **(A)** Immunoblot on lysates from ATP6V1A KO HeLa cells probed for huntingtin, ataxin-2, ataxin-1, and GAPDH. **(B)** Quantification of immunoblot in panel (A), normalized to GAPDH (loading control) levels and to the WT cell line (mean ± SD; analyzed using 2-way ANOVA with post-hoc Šídák’s multiple comparisons tests; **: p<0.01, ns: not significant).

**Figure S4:**
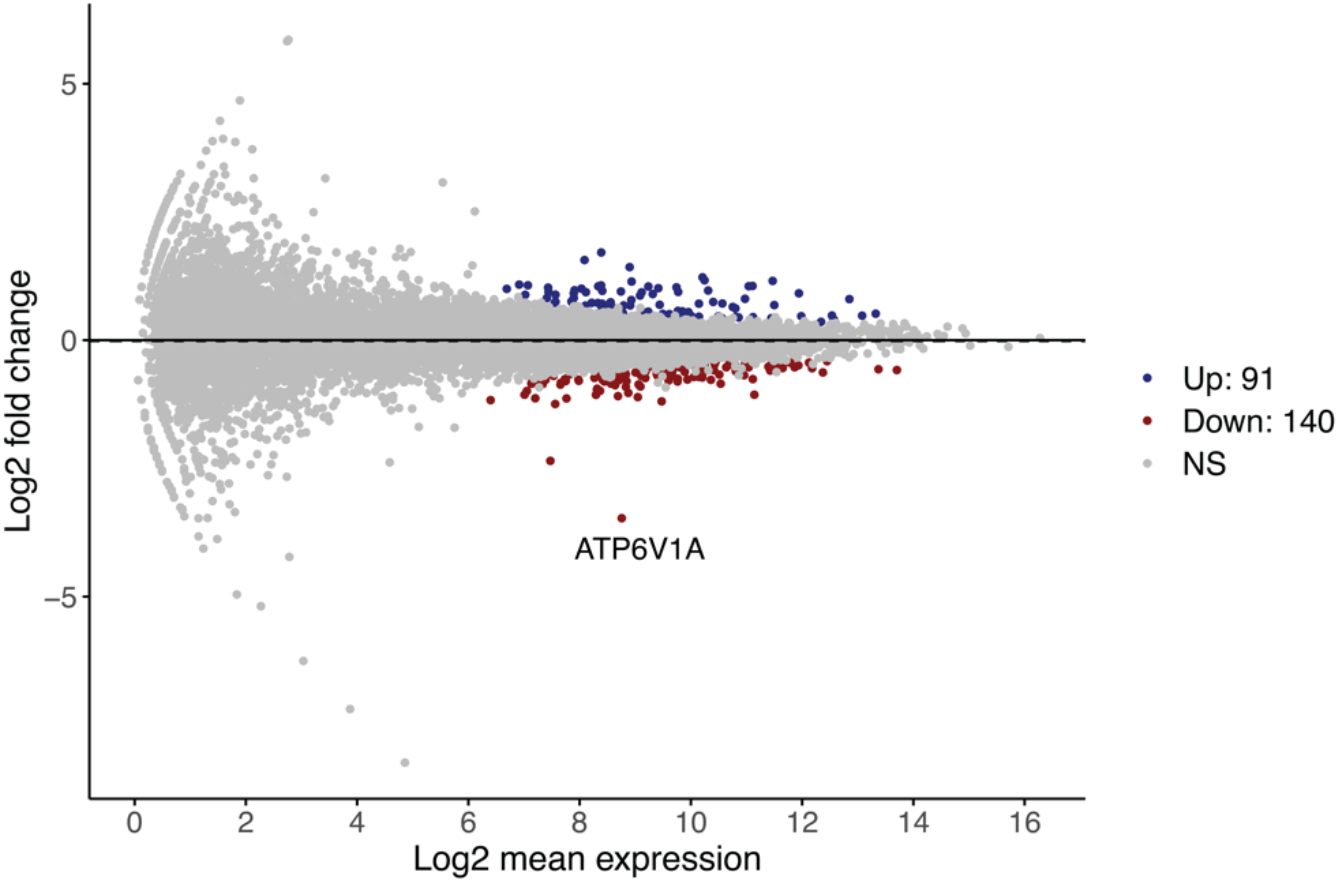
MA plot 72 hours after treatment with NT vs. *ATP6V1A* siRNAs in HeLa cells. To determine whether there are broad transcriptional changes after knocking down a v-ATPase subunit, we performed RNA-seq after HeLa cells were treated with NT or *ATP6V1A* siRNAs. Few noteworthy transcriptional changes are seen (apart from ATP6V1A itself) upon knockdown of *ATP6V1A* (FDR<0.01).

**Figure S5:**
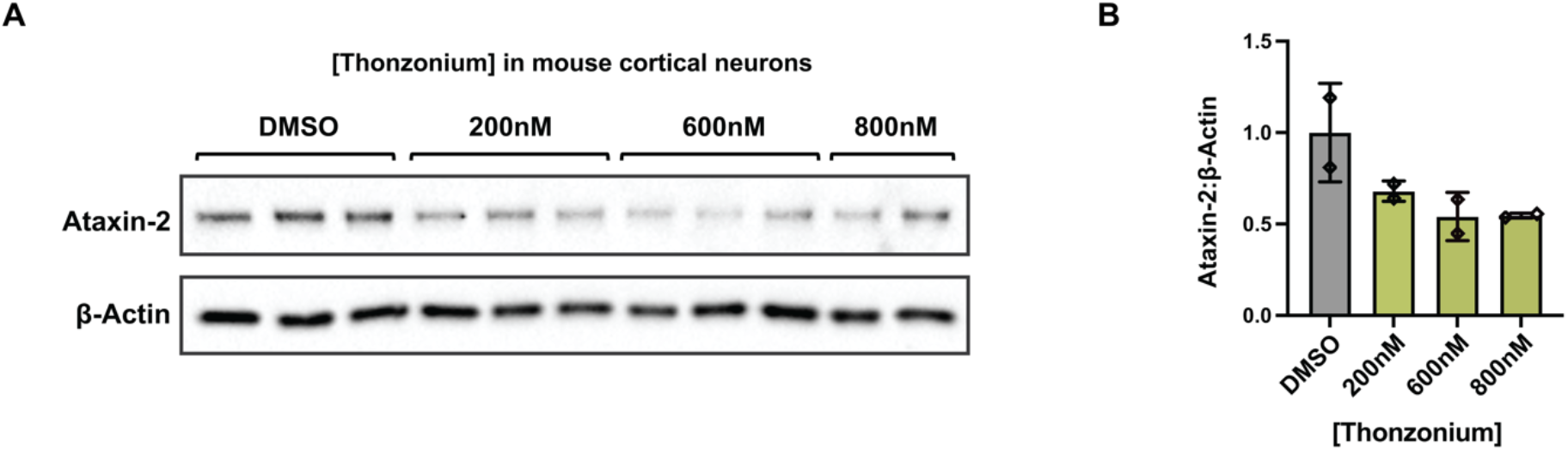
Treating mouse cortical neurons with another drug, Thonzonium, leads to decrease in ataxin-2 protein levels. **a**, Immunoblot on lysates from mouse primary neurons treated with various doses of Thonzonium. **b**, Quantification of the dose-dependent effect of Thonzonium on ataxin-2 (normalized to β-actin) (mean ± SD).

**Table S1: List of hits from screen with β-actin control**. Two biological replicate screens were performed using β-actin as a control in HeLa cells. Hits based on FDR<0.05.

**Table S2: List of hits from screen with GAPDH control**. Two biological replicate screens were performed using GAPDH as a control in HeLa cells. Hits based on FDR<0.05.

## Methods

### Pooled FACS Screen

#### Generating HeLa-Cas9 cells

To generate cells that stably express Cas9, lentivirus containing Cas9 with blasticidin resistance cassette (Cas9-Blast) or blue fluorescent protein (Cas9-BFP) were generated using standard protocols. Low-passage HeLa cells were transfected at 40-50% confluency in a 100 mm dish with lentiviral concentrations such that the infection rate was ∼20%, to reduce the chance that a single cell will incorporate multiple lentiviral particles. 4 days after adding the lentivirus, the culturing media was changed to blasticidin (10 μg/mL)-containing DMEM, high glucose, GlutaMAX™, HEPES media media (Gibco) to select for cells that incorporated Cas9-Blast. The blasticidin-containing DMEM media was replaced every 24 hours until a control plate in parallel of the same quantity of non-Cas9-infected HeLa cells exhibited complete cell death. Cas9-BFP cells were clonally isolated using FACS.

#### Genome-wide deletion library production and titering

All gRNA oligonucleotides were constructed on a microfluidic array, then lentivirus was generated using standard protocols. Briefly, all guides were pooled together at roughly the same concentration (10 sgRNAs per gene targeting ∼21,000 human genes and ∼10,000 safe-targeting sgRNAs), which were cloned into a lentiviral backbone. This pool was used to transfect low-passage HEK293T cells at 70-80% confluency in 150 mm dishes, from which the resulting supernatant contained all 25,000 sgRNAs, with each sgRNA represented ∼1000 times.

#### Generating genome-wide knockout cell line

HeLa-Cas9 cells were cultured in DMEM, high glucose, GlutaMAX™, HEPES media (Gibco) containing 10% FBS (Omega) and 1% penicillin-streptomycin (P/S) (Gibco) in 150 mm plates. The viral media generated above—containing 1000x representation of each sgRNA—was used to infect the HeLa-Cas9 cells. The virus titering was performed such that 5-10% of cells were mCherry-positive, to reduce the chance that a single cell will incorporate multiple gRNAs. 24 hours after infection, media was changed to DMEM media containing puromycin (1 μg/mL) to select for infected cells. The puromycin-containing media was replaced every 24 hours until >90% of cells were mCherry-positive, and an uninfected control plate containing HeLa-Cas9 cells exhibited complete cell death when subjected to puromycin selection. The cells were grown for an additional five days to give Cas9 ample time to cut.

#### Fixation and IF

After trypsinization, approximately 400 million cells of the genome-wide deletion cell line were fixed in 100% methanol for 10 minutes at −20°C. The number of cells to fix was chosen based on ensuring 1000x coverage of the whole genome (250,000 guide elements) while accounting for cells lost during fixation/staining/FACS. After placing in blocking solution (0.4% PBS-Triton containing 5% normal donkey serum and 0.5% BSA) for one hour, primary antibodies against ataxin-2 (1:100; Rabbit; ProteinTech 21776-1-AP) and house-keeping protein GAPDH (1:500; Mouse; Sigma-Aldrich G8795) or β-actin (1:100; Mouse; ThermoFisher Scientific MA1-744) were added to the sample and left overnight at 4°C on a shaker. After rinsing one time in PBS-Triton (0.4%), fluorescent secondary antibodies were added (1:500) for two hours. The cells were then rinsed in PBS, resuspended in PBS containing 2 mM EDTA, and taken directly to the FACS facility for sorting.

#### Fluorescence activated cell sorting

To identify genetic modifiers of ataxin-2 protein levels, the cell suspension was sorted using a BD FACSAria II cell sorter with a 70 μm nozzle (BD Biosciences). Cell populations containing the lowest and highest 20% of ataxin-2 levels—relative to β-actin or GAPDH— were sorted, respectively. Each sorted population, as well as the unsorted (starting) population, were spun down at 600g for 20 minutes at room temperature before extracting genomic DNA.

#### Genomic DNA extraction, PCR amplification, and next-generation sequencing

Genomic DNA was extracted immediately after pelleting using the Blood and Tissue DNeasy Maxi Kit (QIAGEN, 51194). The DNA was isolated according to the manufacturer’s instructions, with the exception of eluting with buffer EB, rather than buffer AE. After PCR amplification using Agilent Herculase II Fusion DNA Polymerase Kit, deep sequencing was performed on an Illumina NextSeq 550 platform to determine library composition. Guide composition between the sorted top 20% and the unsorted (starting) populations were compared using Cas9 high Throughput maximum Likelihood Estimator (casTLE) (Morgens et al., 2016) to determine genes that, when knocked out, increased or decreased ataxin-2 protein levels. Briefly, enrichment of individual guides was calculated as median normalized log ratios of counts between the various conditions. Gene-level effects were then calculated from ten guides targeting each gene, and an effect size estimate was derived for each gene with an associated-likelihood ratio to describe the significance of the gene-level effects. By randomly permutating the targeting elements, the distribution of the log-likelihood ratio was estimated and *P* values derived (Morgens et al., 2016). All data is available under Gene Expression Omnibus accession no. GSE189417.

### Validation & Drug Treatments

#### Cell culture and siRNA transfection

HeLa cells (ATCC^®^) were cultured in DMEM, high glucose, GlutaMAX™, HEPES media (Gibco) containing 10% fetal-bovine serum (FBS) (Omega) and 1% penicillin-streptomycin (P/S) (Gibco) in a controlled humidified incubator at 37°C with 5% CO_2_. For knockdown experiments, cells were reverse transfected with SMARTPool ON-TARGETplus siRNAs (GE Dharmacon) targeting a control siRNA pool (D001810-10) or a gene of interest at a final concentration of 200 nM for 72 hours in 12-well plates, after complexing with DharmafectI (GE Dharmacon) in Opti-MEM (Gibco) for 30 minutes.

SH-SY5Y cells (ATCC^®^) were cultured in DMEM/F12, GlutaMAX™-supplemented media (Gibco) containing 10% FBS (Omega) and 1% P/S (Gibco) at 37°C with 5% CO_2_. siRNA reverse transfection experiments were conducted similarly as for HeLa, except for performing knockdown for 96 hours in two doses (a second dose given after 24 hours with a full media change) and complexing with Lipofectamine RNAiMAX (Invitrogen) in Opti-MEM (Gibco) for 20 minutes prior to addition of cells. Cells were cultured in 24-well plates.

#### Generating ATXN2 mosaic knockout cell line

To generate mosaic *ATXN2* KO HeLa-Cas9 cells, a sgRNA oligonucleotide targeting the first shared exon in *ATXN2* (sequence GATGGCATGGAGCCCGATCC) was cloned into a lentiviral backbone that contains mCherry and a puromycin resistance cassette. This construct was used to transfect low-passage HEK293T cells at 70-80% confluency in 100 mm plates. HeLa-Cas9 cells (cultured in DMEM, high glucose, GlutaMAX™, HEPES media containing 10% FBS and 1% penicillin-streptomycin in 100 mm plates) were then infected with the lentiviral media generated above, such that ∼50% of cells were mCherry-positive. 24 hours after infection, the media was changed to puromycin-containing media (1 μg/mL) to select for cells that received a sgRNA. The puromycin-containing media was replaced every 24 hours until >90% of cells were mCherry-positive, and an uninfected control plate containing HeLa-Cas9 cells exhibited complete cell death upon subjection to puromycin selection. The cells were grown for an additional week to give Cas9 ample time to cut prior to use in flow cytometry.

#### Western Blots

Ice-cold RIPA buffer (Sigma-Aldrich R0278) containing protease inhibitor cocktail (Thermo Fisher 78429) and phosphatase inhibitor (Thermo Fisher 78426) were placed on cells for lysis. After 1-2 minutes, the lysates were moved to Protein LoBind tubes (Eppendorf 02243108), vortexed, and placed on ice. The lysates were vortexed two more times after 10 minute intervals then pelleted at maximum speed on a table-top centrifuge for 15 minutes at 4°C. After moving the supernatant to new Protein LoBind tubes, protein concentrations were determined using bicinchoninic acid (Invitrogen 23225) assays. Samples were denatured at 70°C in LDS sample buffer (Invitrogen NP0008) containing 2.5% 2-mercaptoethanol (Sigma-Aldrich) for 10 minutes. Samples were run on 4–12% Bis–Tris gels (Thermo Fisher) using gel electrophoresis, then wet-transferred (Bio-Rad Mini Trans-Blot Electrophoretic Cell 170-3930) onto 0.45 μm nitrocellulose membranes (Bio-Rad 162-0115) at 100V for 90 minutes. Odyssey Blocking Buffer (LiCOr 927-40010) was applied to membranes for one hour then replaced with Odyssey Blocking Buffer containing antibodies against ataxin-2 (1:1000, ProteinTech 21776-1-AP) and β-actin (1:2000, Thermo Fisher Scientific MA1-744) and placed on a shaker overnight at room temperature. After rinsing three times in PBS-Tween (0.1%) for 10 minutes each, membranes were incubated in Odyssey Blocking Buffer containing HRP-conjugated anti-rabbit IgG (H+L) (1:2000, Life Technologies 31462) or anti-mouse IgG (H+L) (1:2000, Fisher 62-6520) secondary antibodies for one hour. After rinsing the blots three additional times in PBS-Tween (0.1%), the membranes were developed using ECL Prime kit (Invitrogen) and imaged using ChemiDoc XRS+ System and Image Lab software (Bio-Rad Laboratories).

#### RT-qPCR and RNA Quantification

After reverse transfection with siRNAs in 12-well plates as described in the ‘Cell culture and siRNA transfection’ section, the PureLink® RNA Mini Kit was used to isolate RNA with DNase digestion (Thermo Fisher Scientific 12183025). To convert RNA to cDNA, we used the Applied Biosystems High-Capacity cDNA Reverse Transcription kit (Thermo Fisher Scientific 4368813). Each sample had biological triplicates, and technical quadruplicates for each of the replicates. qPCR was performed using TaqMan™ Universal Master Mix II (Thermo Fisher Scientific4440040), using 1 μL of 20X TaqMan gene-specific expression assay to the reaction and our probes of interest (Thermo Fisher Scientific; human *ATXN2*: Hs00268077_m1, human *ACTB:* Hs01060665_g1). The Delta-Delta Ct method was run on the thermocycler and visualized on Thermo Fisher Connect™, from which relative expression values were averaged across all biological/technical replicates per condition.

#### RNA-sequencing

To determine whether there are broad transcriptional changes after knocking down a v-ATPase subunit, we performed RNA-seq after HeLa cells were treated with NT or *ATP6V1A* siRNAs for 72 hours, as described in the ‘Cell culture and siRNA transfection’ section. Briefly, we isolated RNA using the PureLink® RNA Mini Kit with DNase digestion (Thermo Fisher Scientific 12183025), then determined RNA quantity and purity by optical density measurements of OD260/280 and OD230/260 using a NanoDrop spectrophotometer. We assessed structural integrity of the total RNA using a 2200 TapeStation Instrument with RNA ScreenTapes (Agilent Technologies), then prepared mRNA libraries using SureSelect Strand-Specific RNA Library Preparation kit for Illumina (G9691B) on an Agilent Bravo Automated Liquid Handling Platform, following the manufacturer’s protocol. Libraries were sequenced on an Illumina Nova-Seq 6000 machine. Once the data was retrieved, alignment of RNA-sequencing reads to the human hg38 transcriptome was performed using STAR v2.7.3a(Dobin et al., 2013) following ENCODE standard options, read counts were generated using RSEM v1.3.1, and differential expression analysis was performed in R v4.0.2 using the DESeq2 package v1.28.1 (Love et al., 2014) (detailed pipeline v2.1.2 and options available on https://github.com/emc2cube/Bioinformatics/). All data is available under Gene Expression Omnibus accession no. GSE189417.

#### Immunocytochemistry and microscopy

WT or *ATP6V1A* KO HeLa cells were grown on poly-L-lysine-coated glass coverslips [0.1% (wt/vol)] in 24-well plates, then fixed with 4% paraformaldehyde for 30 minutes. Next, the cells were rinsed 3 times with PBS, then blocked with 1% BSA containing 0.4% Triton X-100 for one hour. After overnight primary antibody incubation (1:1000 ataxin-2; 1:1000 β-actin), cells were rinsed 3 times with PBS prior to incubating with fluorescent secondary antibodies (1:500) for one hour. After rinsing with PBS 3 times, coverslips were mounted using Prolong Diamond Antifade mount containing DAPI (Thermo Fisher Scientific). All steps were carried out at room temperature. Images were acquired using a Zeiss LSM 710 confocal microscope. Images were processed and analyzed using ImageJ (version 2.1.0).

#### In vitro drug treatments

Mouse primary neurons were obtained from timed-pregnant, C57BL/6 mice at E16.5 (Charles River). The cortices were dissected out and dissociated into single-cell suspensions with a papain dissociation system (Worthington Biochemical Corporation) and plated onto poly-L-lysine (Sigma Aldrich)-coated plates (0.1% (wt/vol)) at a density of 350,000 cells per well in 24-well plates. The neurons were grown in Neurobasal medium (Gibco) supplemented with P/S (Gibco), GlutaMAX (Invitrogen), and B-27 serum-free supplements (Gibco) at 37°C with 5% CO_2_. After 4 days *in vitro* (DIV), a full media change was performed containing various concentrations of Etidronate (ranging from 1 nM to 100 μM) or water (control), or Thonzonium (ranging from 1 nM to 10 μM) or DMSO (control), and cells were lysed 24 hours later to collect protein. All mouse experiments were approved by the Stanford University Administrative Panel on Animal Care (APLAC).

Human iPSCs-derived neurons (iNeurons) were induced utilizing a Tet-On induction system to express the transcription factor Ngn2. Briefly, iPSCs were maintained in mTeSR1 medium (Stemcell Technologies) on Matrigel-coated plates (Fisher Scientific CB-40230). The following day, doxycycline (2 μg/mL) was added to the media to induce Ngn2 expression, followed by puromycin (2 μg/mL) treatment to rapidly and efficiently induce iNeurons. Three days following induction, the differentiating iNeurons were dissociated using Accutase (STEMCELL Technologies) and resuspended in a culture medium consisting of Neurobasal media (Thermo Fisher), N2 (Thermo Fisher), B-27 (Thermo Fisher), BDNF/GDNF (R&D Systems) on Matrigel-coated assay plates. This resuspension mixture was then plated onto Matrigel-coated 24-well plates. Half-media changes were performed every 2-3 days. 6-7 days after Ngn2 induction, the cells were treated with 10 nM, 100 nM, or 1 μM Etidronate (dissolved in media) or water (control, all to equal volumes).

#### In vivo drug treatments

3-4-month-old WT C57/BL6 mice were given normal (control group) or Etidronate-infused water (10 μM) and MediGel® Sucralose (ClearH_2_O) (30 μM) (treatment group) for voluntary consumption, with 15 animals in each group. After one week, the animals perfused with PBS before dissecting out their brains for flash-freezing. Olfactory bulbs were removed, the hemispheres were separated, and each hemisphere was divided into cortex and cerebellum chunks. Left cortices were then Dounce homogenized and treated for protein extraction in a conventional manner, as described above in the *Western Blots* section.

#### Statistical Methods

Analyses were performed using RStudio (version 1.3.959) or Prism 9.0 (GraphPad), and graphs were generated using Prism 9.0. Data represent mean ± SD. Specific tests (e.g., unpaired t-test, one-way ANOVA, two-way ANOVA, post-hoc tests) and significance are indicated in figure legends.

#### Data and code availability

The data supporting the findings of this study are available from the corresponding author upon reasonable request. All sequencing data is available under Gene Expression Omnibus accession no. GSE189417. Source code for analyzing CRISPR screen data using casTLE method (Morgens et al., 2016) can be found at the following URL: https://bitbucket.org/dmorgens/castle/downloads/. Detailed pipelines and options used for casTLE and RNA-seq are available on https://github.com/emc2cube/Bioinformatics/.

